# Traumatic brain injury modifies adult hippocampal neural stem cell fate to promote neurogenesis at the cost of astrogliogenesis

**DOI:** 10.1101/2023.03.04.531101

**Authors:** P Bielefeld, A Martirosyan, A Apresyan, G Meerhoff, F Pestana, S Poovathingal, N Reijners, W Koning, RA Clement, I Van de Veen, E Toledo, I Durá, S. Hovhannisyan, B. Nilges, A. Bogdoll, N. Kashikar, PJ Lucassen, TG Belgard, JM Encinas, MG Holt, CP Fitzsimons

## Abstract

Moderate Traumatic brain injury (TBI) can result in long-lasting changes in brain function. Although frequently spared from the acute primary injury, the hippocampus becomes affected during a secondary phase that takes place hours, or even days, after TBI, contributing to cognitive deficits. The hippocampus is one of the few brain areas in the adult brain harboring native neural stem cells (NSCs) that continue to generate new neurons (neurogenesis), and to a lesser extent new astrocytes (astrogliogenesis). While deregulation of hippocampal NSCs and neurogenesis have been observed after TBI, very little is known about how TBI may affect hippocampal astrogliogenesis.

Here, we aimed to assess how TBI affects hippocampal NSCs and their subsequent commitment to the neuronal or astroglial lineages. Using a controlled cortical impact model of TBI, single cell RNA sequencing and spatial transcriptomics, we observed a cell population-specific increase in NSC-derived neuronal cells and a decrease in NSC-derived astrocytic cells. These cellular changes were associated with cell-population specific changes in gene expression and dysplasia within the dentate gyrus.

Overall, our findings support the conclusion that TBI modifies adult hippocampal NSC fate to promote neurogenesis at the cost of astrogliogenesis, and highlights specific cell populations as possible targets to counteract the changes induced by TBI in the hippocampus.

## Introduction

Traumatic brain injury (TBI) is a major global health problem linked to everyday life events such as domestic activities, participation in (contact) sports, road accidents and occupational hazards, in which head trauma causes brain damage (PMID: 35259824). 69 million people worldwide suffer from TBI annually (PMID 29701556). While TBI-induced brain damage is the leading cause of death below 45 years of age (PMID: 26903824), nearly half of milder TBI patients experience some form of long-term cognitive impairment (PMID 24529420) and are more likely to develop depressive symptoms and neurodegenerative diseases (PMID: 29355429; PMID: 29190146; PMID: 36759368). Despite these devastating consequences for patients, there is currently a lack of specific therapeutic targets, or effective drug treatments, for TBI (PMID: 35662186, PMID: 35624914).

Although frequently spared from the acute primary injury, the hippocampus generally becomes affected during a secondary injury phase, which spreads throughout the brain after the initial TBI (PMID: 26903824). Importantly, increasing evidence indicates that cognitive dysfunction after TBI is associated with changes in hippocampal function (PMID: 26903824), that occur during this secondary phase. The hippocampus is critical for cognition, and also one of the few areas in the adult brain that harbors native neural stem cells (NSCs), that have been implicated in cognitive and emotional control (PMID: 29679070). Upon activation, NSCs generate proliferative progenitor cells and neuroblasts, which give rise to immature dentate granule neurons (PMID: 16702546). In addition, NSCs generate new astrocytes (astrogliogenesis) under physiological conditions (pmid 26729510, 21549330, PMID: 35451150, PMID: 35794479). These newly generated neurons and astrocytes persist in a specialized anatomical location in the subgranular zone of the Dentate Gyrus (DG), termed the adult hippocampus neurogenesis (AHN) niche (PMID: 10975875, PMID: 26330519, PMID: 10975875). Although the functional contribution of some of the NSC-derived neuronal cell types within the AHN niche has been studied (PMID:24090877, PMID: 16144763), the properties of the adult NSC-derived astrocytes remain largely uncharacterized (PMID: 35451150).

Here, we aimed to assess in detail how TBI affects NSC fate in the adult hippocampus, leading to changes in neurogenesis and astrogliogenesis and, consequently, in the relative cellular composition of the AHN niche. We applied a controlled cortical impact model of TBI (PMID: 35618831) to a transgenic reporter mouse line in which GFP is expressed in individual cells of the AHN niche (PMID: 14730584; PMID: 31222184), under the control of the neuroepithelial stem cell protein (Nestin) promoter. The expression of GFP in this mouse line has been previously used in single cell RNA sequencing (scRNA-seq) studies of the AHN niche to sort NSCs, neural progenitor and other cell populations, as well as exclude that populations of NSCs are dominated by astrocytes, due to the otherwise close similarity between the two cell types (PMID: 26299571; PMID: 29241552, PMID: 29241552, PMID: 33581058). Using scRNA-seq in combination with spatial transcriptomics technology, we show that TBI disturbs the balance between NSC-driven neuro- and astrogliogenesis, effectively reducing the numbers of NSC-derived astrocytes, while increasing the numbers of NSC-derived neuronal cells. In addition, we molecularly characterize several novel cell populations derived from hippocampal NSCs. Finally, we trace back these cell populations *in situ*, uncovering significant changes in the anatomical location of NSC-derived cell populations in the DG after TBI. As such, our work provides a basis for future investigations of specific cell populations that could serve as targets to counteract the changes induced by TBI in the hippocampus (PMID: 30254269), and may help us to better understand the role of NSCs in hippocampus-dependent cognition.

## Results

### TBI induced impairments in hippocampus-dependent cognition correlates with cellular changes in the dentate gyrus

First, we characterized cellular changes in the hippocampi of mice subjected to mild-moderate TBI and how this correlates to changes in hippocampus-dependent cognition. For this, we compared sham craniotomized mice (Control) and mice subjected to unilateral controlled cortical impact (TBI). 15 days post-surgery, TBI was found to have induced unilateral hippocampal astrogliosis, measured as increased GFAP+ cell coverage area (Fig. 1a-d), and increased neurogenesis as assessed by the numbers of DCX+ cells in the DG (Fig. 1e-g). DCX+ cells could be further subdivided into 6 categories, according to the presence and shape of their apical dendrites: A, no processes; B, stubby processes; C, short horizontal processes; D, short vertical processes oriented to the molecular layer; E, one long vertical dendrite; F, long branched vertical dendritic tree, as described in (PMID: 17105671). We found a significant increase in category C and D cells (Fig. 1h), which have been classified largely as early NBs that express the proliferation marker *Mki67* (PMID: 17105671). These cellular changes correlated with impaired performance in the Morris water maze, a commonly used test for hippocampal learning and memory. Mice subjected to TBI showed significant deficits in learning the position of a hidden platform in the test pool (Fig. 1i, j). Together these results suggest that TBI induced impairments in cognition result from changes in the cellular architecture of the hippocampus.

**Figure 1.**
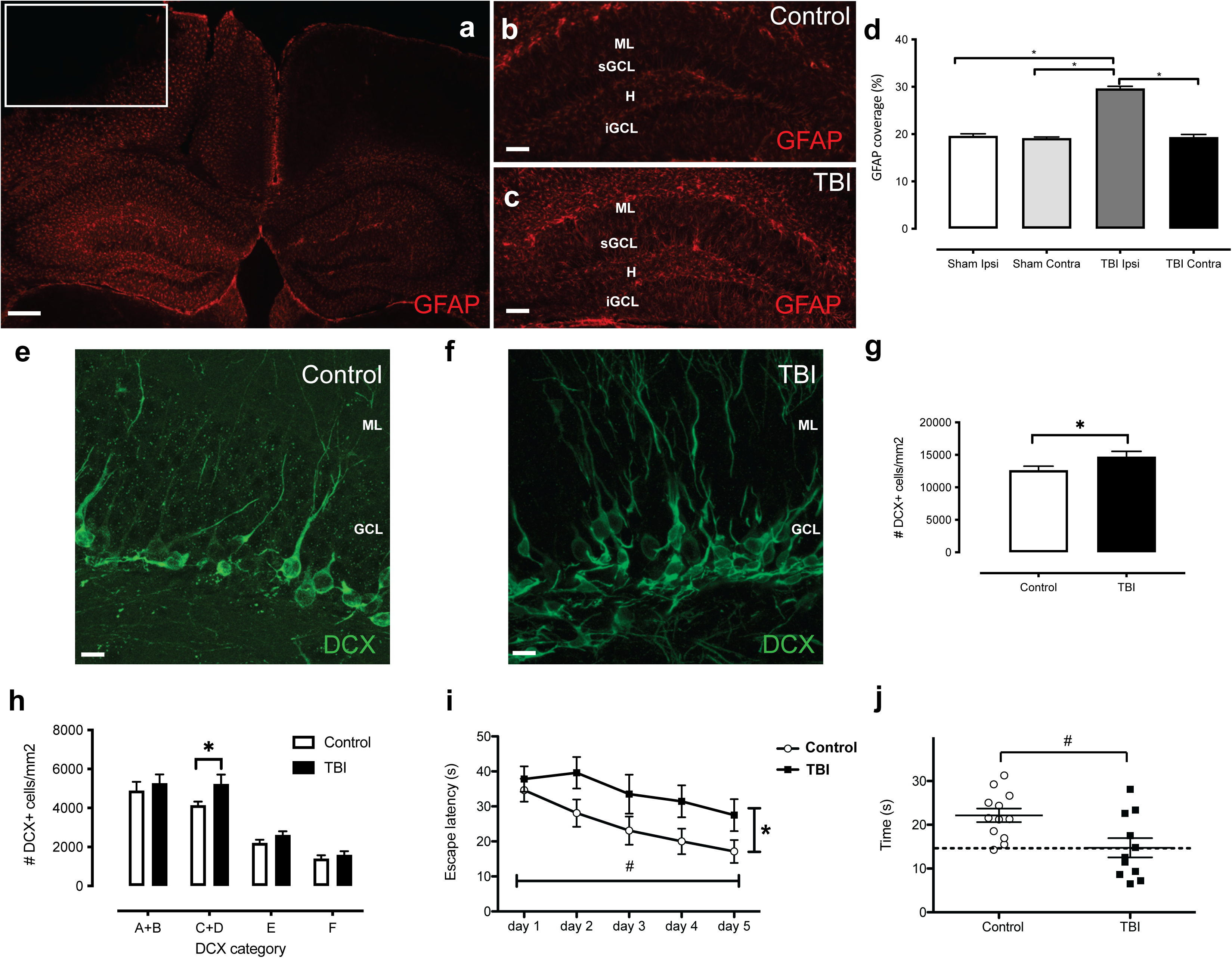
Immunohistochemical and behavioral characterization of hippocampal changes following TBI. Representative example of GFAP (red) immunohistochemistry showing ipsilateral astrogliosis (a), white box indicates the location of the controlled cortical impact, scale bar, 250 µm. Higher magnification examples of increased astrogliosis in the hippocampus of Control (b) vs. TBI (c) mice; ML: molecular layer, sGCL: suprapyramidal granule cell layer, H: hilus, iGCL: infrapyramidal granule cell layer. Scale bars, 100 µm. Quantification of GFAP surface coverage in the dentate gyrus (n=3), *p<0.05 one way ANOVA with Tukey’s post-hoc test (d). High magnification images showing DCX+ cells in the DG of Control (e) vs. TBI (f) mice. Scale bars, 25 µm; ML: molecular layer, GCL: granule cell layer. Quantification of DCX+ cells in the dentate gyrus (g), (Control n=5, TBI n=4, *p<0.05 unpaired t-test). Quantification of DCX+ cell phenotypes according to the presence, shape and orientation of apical dendrites (PMID: 1710567) within the dentate gyrus (h), (Control n=5, TBI n=4, *p<0,05 one way ANOVA with Tukey’s post-hoc test). Escape latency in the Morris Water Maze test (i), (Control n=12, TBI n=11; #p<0.05 time, *p<0.05 treatment, 2-way repeated measures ANOVA with Bonferroni post-hoc test. Percentage of time spent in target quadrant during Morris Water Maze probe trial (j), (Control n=12, TBI n=11, #p<0,05, unpaired t-test).

### A single cell census of the AHN niche reveals that cell identity is maintained after TBI

To understand the cellular changes induced by TBI in DG NSC and their progeny in a comprehensive but unbiased manner we used scRNA-seq. Nestin-GFP mice were divided randomly into Control and TBI groups. Fifteen days post-surgery, DGs were micro-dissected, dissociated and single Nesin-GFP+ cells isolated using FACS (in combination with a live/dead dye to eliminate dead cells) (Fig S1). Cells were then pooled and subjected to scRNA-seq using the 10X 3’ whole transcriptome analysis workflow. An unbiased integration and clustering approach was then applied to the data set (which contained 7791 high-quality cells), using the Seurat algorithm (PMID: 31178118). We were able to identify 10 cell clusters representing the expected major cell types of the DG (Fig. 2a, Table S1). Based on the expression levels of known marker genes, we defined abundant cell clusters containing NSCs (*Neurog2^+^, Hmgn2^+^, Sox4^+^, Sox11^+^, Mki67^+^),* Radial Glia-like (RG-like) cells (*Ascl1^+^*, *Ccnd2^+^, Vim^+^, Hes5^+^, Mki67^−^),* astrocyte-like cells (*Slc1a3^+^, Aqp4^+^, S100b^+^, Aldoc^+^*), neuronal-like cells (*Neurod1^+^*, *Snap25^+^, Dcx^+^*), and oligodendrocytes (Oligo) (*Mog^+^, Mag^+^, Mbp^+^*). In addition, other less abundant cell populations were identified, such as oligodendrocyte precursor cells (OPCs) (*Pdgfra^+^, Gpr17^+^, Mag^+^*), pericytes/mural cells (*Des^+^, Col1a2^+^*), endothelial cells (*Pecam1^+^, Flt1^+^*) and microglia (Mgl^+^, *CD68^+^, Cx3CR1^+^*). Crucially, UMAP representation indicated that TBI does not seem to induce significant changes in the overall cellular identity of the DG (Fig. 2b), although significant alterations in the proportions of various cell types present were detected (Fig. 2c, Table S1).

**Figure 2.**
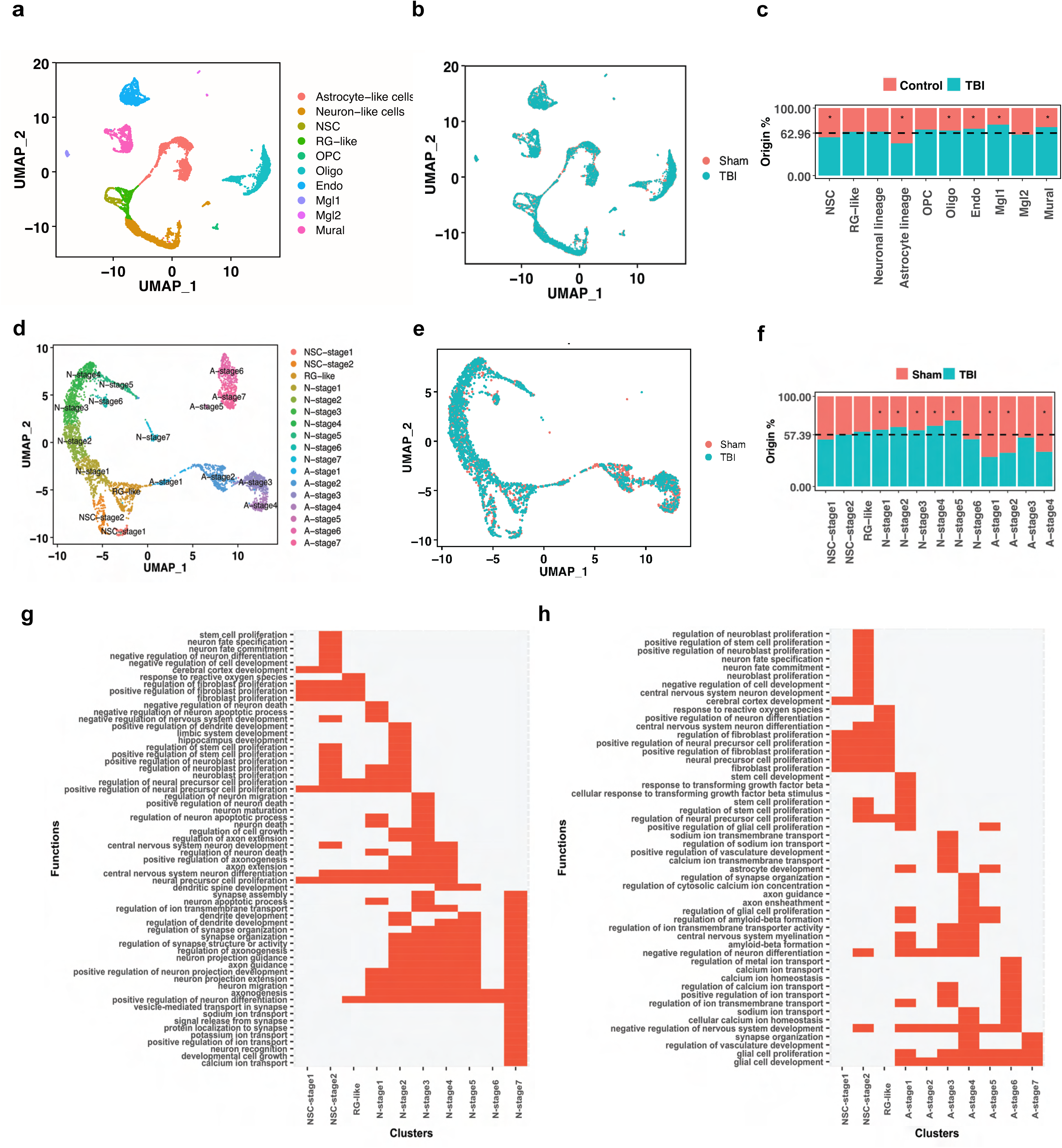
Molecular characterization of the NSC-derived neuro- and astrogliogenic lineages within the mouse dentate gyrus and the effects of TBI. UMAP-based visualization of the major higher-order cell types identified by Seurat in the single cell dataset. Each dot represents a single cell. Cells with similar molecular profiles group together. Cell types were assigned according to the expression of specific marker genes and are labelled in different colors (a). UMAP-based visualization of single cell data according to experimental origin (Control or TBI) (b). Bar plots showing the relative number of cells per cluster originating from Control or TBI samples against their predicted abundance (based on 62.96% of all cells originating from TBI samples: dashed line), *p<0.05 vs. expected abundance, binomial test (c). UMAP-based visualization of NSC-derived neuronal and astrocytic lineages obtained by extraction of neuronal and astrocytic cells followed by reclustering. Each dot represents an individual cell and discrete cell states are identified in specific colors (d). UMAP-based visualization of reclustered data according to experimental origin (Control or TBI) (e). Bar plots showing the relative number of cells per identified cell state in Control or TBI samples, against their expected abundance (based on 57.39% of all cells originating from TBI samples: dashed line), *p<0.05 vs. expected abundance, binomial test (f). GO biological pathway matrix for the cell clusters included in the neuronal (g) and astrocytic (h) lineages.

Based on the developmental origin of neurons and astrocytes from radial glia-like NSCs in the hippocampus (PMID: 33349709, PMID: 35451150), and our own observation of alterations in both astrocytes and neurons following TBI (Fig. 1), we decided to examine the effects of TBI on neurogenesis and astrogliogenesis in greater detail. For this, the NSC, RG-like, astrocyte-like and neuronal-like populations were extracted and re-clustering was performed. This approach revealed several previously unidentified populations (Fig. 2d, Table S2), including 3 clusters with gene expression profiles characteristic of NSCs. These three NSC clusters expressed known NSC markers (*Ascl1, Vim, Hes5, Id4*) (Fig. S2), and could be further differentiated into 3 subgroups, based on marker expression: NSC-stage 1 (expressing *Ranbp1, Ezh2, Nme1*), NSC-stage 2 (expressing *Eif4g2, Hspa5*) and RG-like (expressing *Neurog2, Zeb 1*) (Fig. 2d; Fig. S3; Table S3). Importantly, NSC-stage 1 and 2 cells robustly expressed the proliferation marker *Mki67,* but only few RG-like cells did (Fig. S3; Table S3). To investigate the neurogenic and astrogliogenic pathways in more detail, we performed a pseudotime lineage analysis on the data using the Monocle algorithm (PMID: 30787437), with the identified NSC populations as the developmental root. Using this methodology, neuronal populations could be divided in UMAP space into a neuronal lineage which included NSC-stage 1 and 2, RG-like cells and other clusters that expressed *Dcx* and *NeuroD1, Mdk, Prox1 (*N-stage 1-2 cells); *Plxna4, NeuroD2, Prox1* (N-stage 3-4 cells); *Tubb2b, Fez1, Prox1* (N-stage 5-6, cells) (Fig S3 and Table S3). N-stage 7 cells, expressing *Syt1 and Reln* clustered independently from N-stage 1-6 cells in UMAP space, indicating they likely belong to a different lineage not derived from NSCs (Fig. 2d). Pseudotime analysis also revealed an astrocytic lineage including NSC-stage 1 and 2, RG-like and four astrocytic cell clusters dubbed A-stage 1-4 cells (Fig S3 and Table S3). A-stage 1 cells expressed NSC markers (*Ascl1, Vim, Hes5*) and genes commonly expressed in astrocytic cells (*Gfap, Ntrk2, id2*), suggesting A-stage 1 cells are immature astrocytic cells originating from NSCs. A-stage 2 cells did not express the NSC markers *Ascl1, Vim and Hes5*, but expressed the astrocytic markers *Gfap, Ntrk2 and id2,* presumably representing cells differentiating along the astrocytic lineage. A-stage 3-4 cells expressed *Gfap* but differed from A-stage 2 cells by expressing a different set of astrocytic markers, including *Smo, Fgfr3, Dmd* and *Hes5*. A-stage 5-7 cells clustered separately in UMAP space, as they neither expressed NSC markers (*Ascl1, Hes5*) nor the astrocytic markers *Smo, Fgfr3 and Dmd*. They did, however, express *Gfap, Ntrk2 and id2*, indicating that they likely represent a different astrocyte linage that is not derived from NSCs (Fig. 2d). Interestingly, TBI does not seem to induce significant changes in the overall developmental trajectories of NSC-derived neurons and astrocytes (Fig. 2e, f). However, several NSC-derived neuronal cell clusters (N-stage 1, 2, 4, 5) were enriched in the cell suspensions prepared from animals subjected to TBI, with N-stage 5 cells showing the largest TBI-induced effect (Fig. 2f, Table S2). In addition, several NSC-derived astrocytic cell clusters were depleted in these cell suspensions, with A-stage 1, 2 and 4 cells showing a significant TBI-induced reduction, while A-stage 3 cells were unaffected (Fig. 2f, Table S2).

Gene ontology (GO) biological pathway (BP) analysis performed for the neuro- and astrogliogenic lineages confirmed 2 trajectories, with distinct changes in associated biological functions over pseudotime, indicating a gradual change towards neuronal (Fig. 2g, Table S4, Table S5) or glial (Fig. 2h, Table S4, Table S5) functions, respectively. Interestingly, GO analysis suggests NSC-stage 2 is the cell population with the highest proliferative potential, suggesting these cells represent activated NSCs, in agreement with their robust *Mki67* expression (Fig. S2, Fig. S3, Table S3). Additionally, GO pathway analysis indicated that several astrocytic cell clusters may retain some proliferative potential, even in the absence of active proliferation, indicated by their lack of *Mki67*expression (Fig. 2h). In the absence of major effects of TBI on lineage and cell type identity, the change in the relative proportions of the various neuronal and astrocyte subgroups suggests that TBI affects principally cell fate determination.

### RNA velocity predicts NSC fate changes after TBI

RNA Velocity is a computational tool for predicting future cell state from scRNA-seq data, by analyzing the ratio of spliced versus unspliced RNA transcripts (PMID: 30089906; PMID: 32747759). RNA velocity analysis performed on our scRNA-seq dataset confirmed our basic finding of neuronal and astrocytic lineages originating from NSCs in the AHN (Fig. 3a, b). Based on the calculated transition probabilities, it appears that both NSC-stage-1 and NSC-stage-2 cells transition to RG-like cells (Fig. 3c-h), which in turn feed both the neuronal (N-stage 1) and astrocytic (A-stage 1) lineages (Fig. 3i-q). Crucially, in Control animals, the probability of RG-like cells to transition to astrocytic lineage is higher than the probability of transitioning to the neuronal lineage (Fig. 3k). Consistent with the numerical changes observed in specific cell clusters after TBI (Fig. 2f), RNA velocity analysis suggests TBI promotes an increase in the number of cells entering the neuronal lineage by promoting the transition of NSC-stage 1 cells to NSC-stage 2 (Fig. 3e), as well as promoting the transition of both NSC-stage-2 and RG-like cells to N-stage 1 (Fig. 3h, k). In contrast, the probability of RG-like cells progressing to A-stage 1 reduced (Fig. 3k), suggesting that increased neurogenesis occurs at the expense of astrogliogenesis. Interestingly, in Control conditions, a proportion of NSC-stage-2 cells can transition to N-stage-1, an effect promoted in TBI conditions (Fig. 3 h), further tipping the balance of cell production towards neurogenesis. Consistent with an imbalance in cell numbers, TBI seems to have a strong effect on the apparent stable identity (self-renewal) behaviors of both NSC-stage-1 and RG-like cells (Fig. 3e, k). In summary, therefore, our RNA velocity analysis is entirely consistent with the major effect of TBI being on cell fate decisions rather than cell identity *per se*.

**Figure 3.**
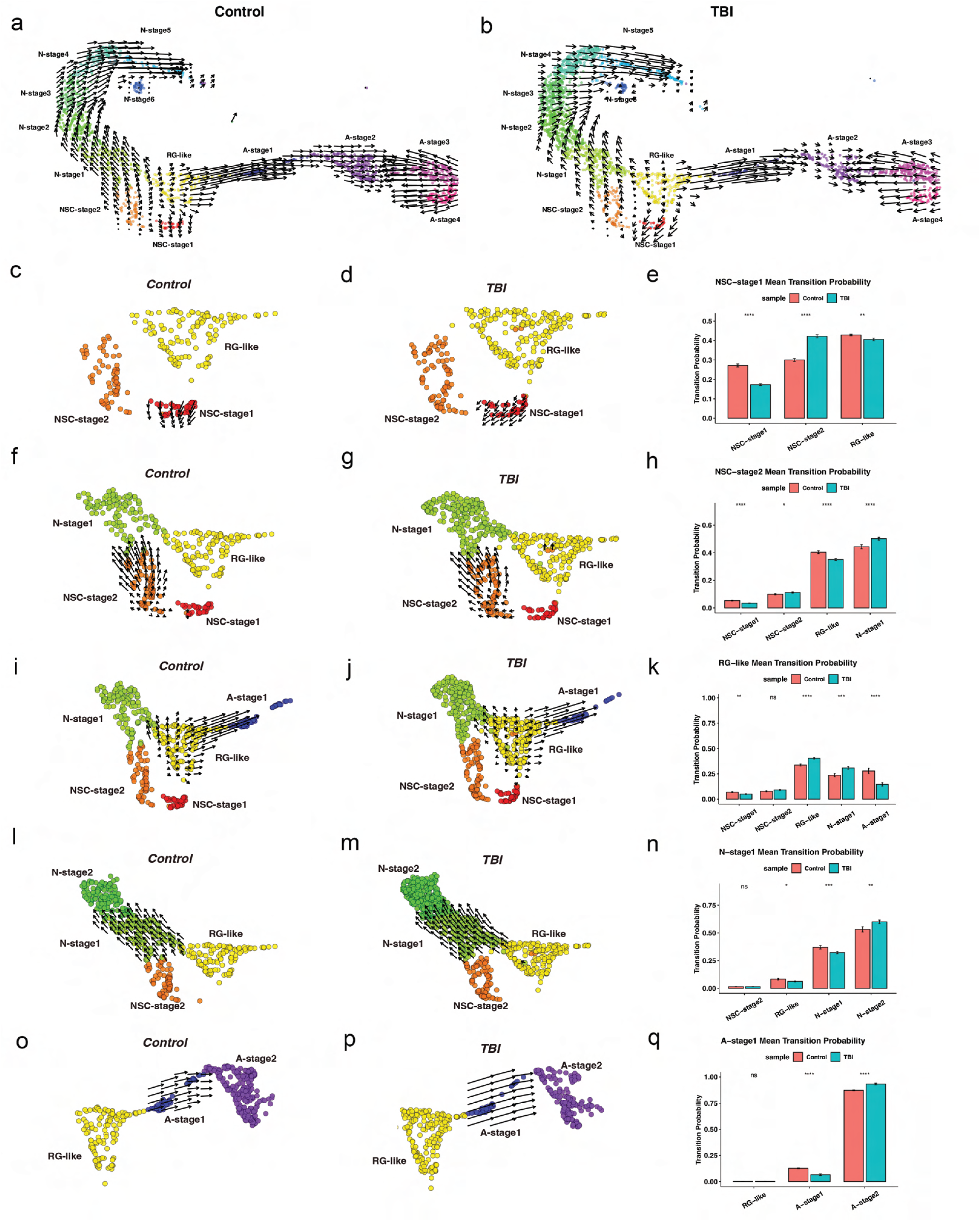
RNA velocity analysis indicates that TBI induces a subtle but functionally significant shift in hippocampal neural stem cell fate. RNA velocity analysis along both the neuronal and astrocytic lineage in the Control (a) and TBI (b) groups. Left-to-right, RNA velocity analysis on Control (left) and TBI (middle) cells with mean transition probabilities indicated (right) NSC-stage 1 cells (c, d, e), NSC-stage 2 cells (f, g, h), RG-like cells (i, j, k), neuronal N-stage 1 cells (l, m, n), and astrocytic A-stage 1 cells (o, p, q). In all panels, vectors (arrows) indicate the predicted direction and speed of movement of single cells in transcriptome space, colors indicate previously characterized cell clusters indicated in the figures; ns: p > 0.05; *p<0,05; **p <= 0.01; ***p <= 0.001; ****p<= 0.0001, independent non-parametric Wilcoxon test.

### Differential gene expression analysis reveals cell population-specific responses to TBI

Next, we investigated the cell-type specific alterations in gene expression induced by TBI. For this, we compared gene expression patterns per cell population between Control and TBI groups using the Wilcoxon Ranked Sum test. We found several differentially expressed genes (DEGs) in specific cell populations. Overall, TBI resulted in more upregulated genes (38 in total across 9 cell populations) than downregulated genes (16 in total across 8 cell populations) (Fig. 4a, b; Table S6). A-stage 3 was the population with the largest number of DEGs, with 11 upregulated and 5 downregulated genes, followed by N-stage 4 with 7 upregulated and 5 downregulated genes and A-stage 2 with 10 upregulated and 1 downregulated gene. TBI did not, however, induce large effects on gene expression in NSC populations. In RG-like cells, two genes were found upregulated: Mediator Complex Subunit 29 (*Med29*) and protein phosphatase 1 regulatory inhibitor subunit 14B (*Ppp1r14b*). The Mediator Complex is a large multiprotein coactivator essential for both activated and basal transcription (PMID: 14576168) and *Med29* is amplified and overexpressed in hyperproliferative cells (PMID: 17332321, PMID: 21225629), suggesting it may be involved in transcriptional and proliferative changes associated with fate changes in RG-like cells (consistent with RNA velocity predictions). A detailed examination of DEGs across cell populations, revealed that *Ppp1r14b*, a gene linked to cell proliferation, growth and apoptosis (PMID: 35679681, 34858479), is consistently upregulated by TBI in 7 cell populations (Fig. 4a, Table S6). Specifically, *Ppp1r14b* is upregulated by TBI in RG-like (Fig. 4c), NSC-stage 2, (Fig. 4d), astrocytic A-stage 3 (Fig. 4e), and neuronal N-stage 1-4, (Fig. 4f-i) cell populations, suggesting that *Ppp1r14b* may play an important role in the coordinated action of TBI across these cell populations. GO pathway analysis of the DEGs between Control and TBI groups in specific populations indicated that several biological processes (BPs) were affected by TBI (Fig. S4, Table S7, Table S8). Specifically, several proliferation-associated BPs were affected in NSCs and derived neuronal populations (NSC-stage 1, NSC-stage 2, N-stage 2 and N-stage 3 cells) (Fig. S4). In the NSC-derived astrocytic populations, several BP associated with apoptosis, DNA damage, transcription, ribosome assembly and biogenesis, and microtubule polymerization were affected by TBI (Fig. S4).

**Figure 4.**
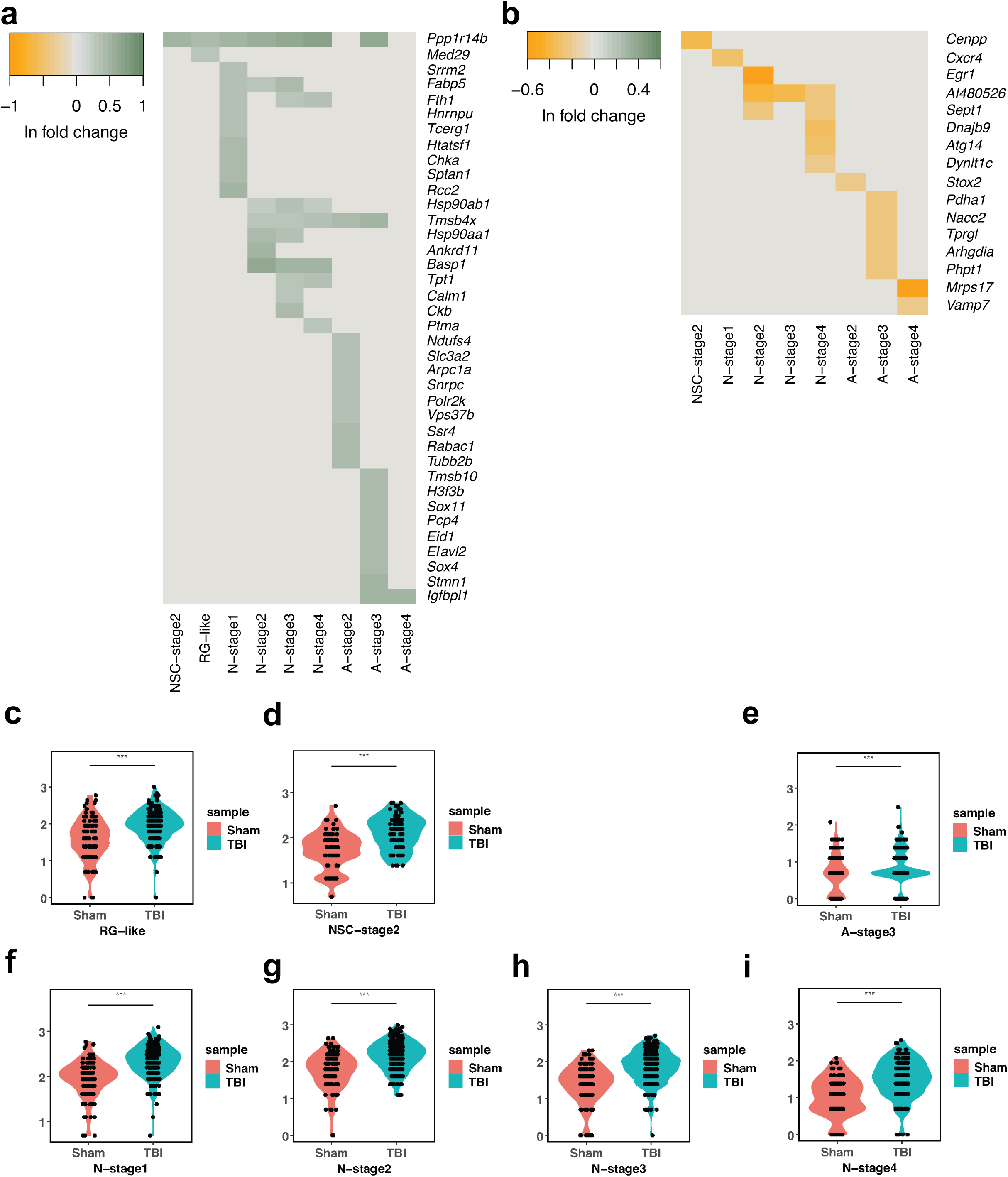
TBI induces cell population-specific changes in gene expression in NSCs and NSC-derived cells in the DG. Heatmaps showing expression of individually upregulated (a) and downregulated (b) genes across NSCs and NSC-derived cell populations. Color bars indicate relative intensity of expression, indicated as ln fold-change in TBI vs. Control groups; SCT normalized data are compared. Violin plots showing changes in *Ppp1r14b* expression induced by TBI in RG-like cells (c), NSC-stage 2 cells (d) astrocytic A-stage 3 cells (e) and neurogenic N-stage 1-4 cells (f-i). ***p<0.01, MAST test with Bonferroni *post-hoc* test on SCT normalized data.

### TBI affects the anatomical location of specific NSC-derived populations in the DG

Our scRNA-seq results indicate the presence of previously uncharacterized cell populations in the DG. To validate their existence, we performed multiplexed fluorescence *in situ* hybridization (FISH) using RNAscope (PMID: 27339989), in hippocampal slices of wild type mice, focusing on the identification of NSC-derived astrocytic cells, for which we could find a suitable set of probes, including *Slc1a3, Hapln1, FrzB, Ascl1, Sparc, Sned1* and *Neat1* (Fig. S5). We were able to localize three of the astrocytic cell populations in the DG identified in our scRNA-seq experiments: A-stage 4 cells (*Slc1a3+, Hapln1+, Neat1+, Sned1+, Sparc-, Ascl1-, Frzb-*; Fig. 5a), A-stage 2 cells (*Slc1a3+, Hapln1-, Neat1+, Sned1+, Sparc-/+, Ascl1-, Frzb-*; Fig. 5b) and A-stage 1 cells (*Slc1a3+, Hapln1-, FrzB+, Ascl1+, Sparc+, Neat1+, Sned1+,* Fig. 5c). These different cell populations were observed in distinct locations within the DG. A-stage 1 and 2 cells, were located in the subgranular zone (SGZ) (Fig. 5d, insets d’ and d’’, respectively), while A-stage 4 cells were located in the molecular layer (ML), close to the external layers of the granule cell layer (GCL) (Fig. 5d, inset d’’’). Indeed, NSCs and their neuronal and astrocytic progenies are found in SGZ, and this anatomical positioning has been linked to their functional roles and integration into hippocampal circuits (PMID:24090877, PMID: 35451150, PMID: 31346164). Overall, our results with RNAscope confirmed that early NSC-derived astrocytic cells (A-stage 1 and 2) locate to the SGZ, while other populations in the NSC-derived astrocytic lineage may reside in more distant locations. Unfortunately, however, RNAscope is restricted to the simultaneous detection of 12 probes. To overcome this limitation, we aimed to characterize the anatomical location of NSCs and NSC-derived neurogenic and astrocytogenic cells using the recently developed Molecular Cartography spatial transcriptomics platform (Resolve Biosciences GmbH, Monheim am Rhein, Germany) (PMID: 35021063). Molecular Cartography allows for simultaneous detection of up to 100 transcripts. We designed an extended panel of 93 probes to localize cell populations identified from the scRNA-seq dataset in the DG of mice from Control and TBI groups (Fig. S5). In a first validation step, the localization of probes in Control tissue was compared to the corresponding signals obtained from the Allen Brain Atlas (PMID: 17151600), which showed significant overlap (Fig. S5, Table S9). Using the multimodal reference mapping tool of the Seurat package, we then compared the signals from the 93 Molecular Cartography targets to the scRNA-seq dataset. This analysis indicated a substantial overlap in the cell populations identified by both techniques (Fig. 5g, h), with all NSC and NSC-derived populations defined in the scRNA-seq data found back in the Molecular Cartography signal clusters (MCCs) (Fig. 5h). Supporting the high concordance between both techniques, pseudotime analysis of the merged scRNA/MCCs dataset confirmed the presence of NSC-derived neurogenic and astrocytogenic lineages (Fig. 5i, j). The relationship between the two datasets was not always one-to-one, as some of the populations defined by scRNA-seq were represented in more than one MCC (Fig. 5k). However, the high degree of overlap between the datasets indicates that Molecular Cartography allows the spatial localization of scRNA-seq defined NSC and NSC-derived cell populations in the intact DG.

**Figure 5.**
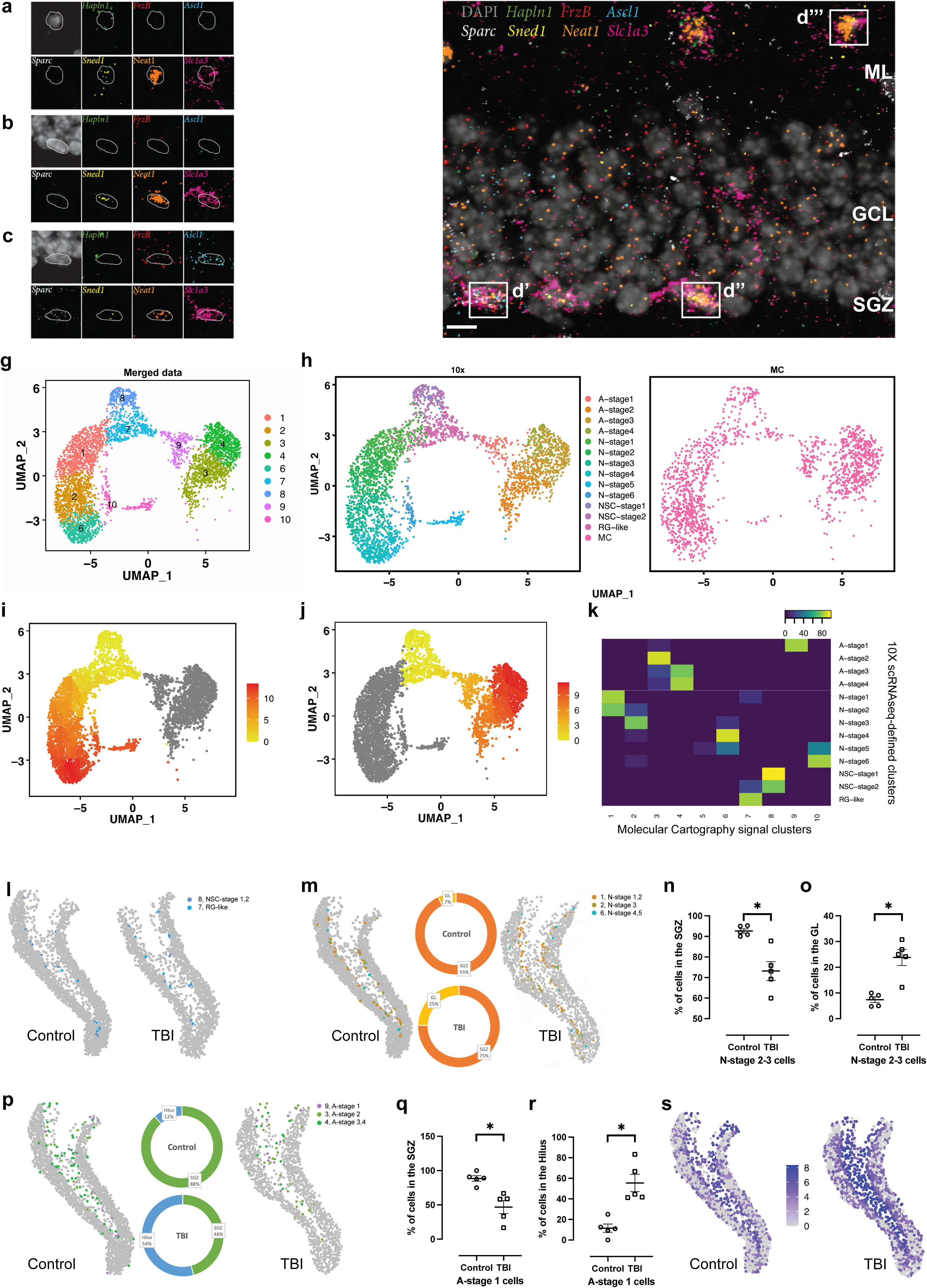
TBI induces changes in the location of specific cell populations in the dentate gyrus. RNAscope visualization of marker genes used to identify individual A-stage 4 (a), A-stage 2 (b) and A-stage 1 (c) cells in intact tissue. Overview of the intact dentate gyrus (d) with individual A-stage 4 (d’’’), A-stage 2 (d’’) and A-stage 1 (d’) cells illustrated, ML: molecular layer, GCL: granule cell layer, SGZ: subgranular zone; scale bar, 5 µm. UMAP-based representation of the combined 10X and Molecular Cartography (MC) datasets, with colors identifying the 10 distinct cell states identified in the MC dataset (g). Both scRNA-seq derived (left) and MC-derived data (right) share a similar distribution within UMAP space, with colors in the 10X dataset indicating neuronal and astrocytic cell lineages identified in the original scRNA-seq data (h). UMAP-based representation of pseudotime for both neuronal (i) and astrocytic (j) lineages using the combined 10X-MC dataset, with color bars indicating relative pseudotime distance from the root population (yellow: minimum, red: maximum). Correlation matrix showing the relative overlap in gene expression data between cell populations identified by scRNA-seq (10X scRNA-seq-defined clusters) and Molecular Cartography (k). Color bar indicates the % of genes in a scRNA-seq-identified cluster present in a cluster identified by Molecular Cartography. Spatial mapping of NSC populations in Control and TBI groups in the DG (l), with colored dots indicating the location of single cells belonging to the populations indicated in the legend. Spatial mapping of NSC-derived neuronal populations in Control and TBI groups in the DG (m), with colored dots indicating the location of single cells belonging to the populations indicated in the legend; pie charts represent the % localization of N-stage 2-3 cells to the SGZ (orange) or the GL (yellow). Bar graphs showing the quantification of N-stage 2-3 cells in the SGZ (n) or the GL (o) in Control and TBI groups. Spatial mapping of NSC-derived astrocytic populations in Control and TBI groups (p), colored dots indicate the location of single cells belonging to the populations indicated in the legend; pie charts represent the % localization of astrocytic A-stage 1 cells to the SGZ (green) or the Hilus (blue). Bar graphs showing the quantification of A-stage 1 cells in the SGZ (q) or the Hilus (r) in Control and TBI groups, *p<0,05, unpaired t-test. Spatial mapping of *Gfap+* astrocytes in the dentate gyrus from Control and TBI animals (s); dots indicate the location of single cells, with color encoding the relative intensity of *Gfap* expression. Spatial representations in l, m and p are representative examples of hippocampal slices analyzed to generate the data in n, o, q, and r.

We then asked whether TBI affects the location of specific cell populations within the DG. First, we segmented the DG into 4 different regions: SGZ, inner GCL (GL1); mid and outer GCL (GL2), and hilus, as described in (PMID: 12466205). Then, we compared the location of NSCs and the corresponding neuronal and astrocytic lineages in the Control and TBI groups, using the MCCs defined in Fig. 6k. We were unable to detect any change induced by TBI in the location of cells in MCCs 7 and 8 (RG-like, NSC-stage 1 and NSC-stage 2 cells) (Fig. 5l). However, the location of cells in MCC 2 (N-stage 2-3 cells) and MCC 9 (A-stage 1 cells) was affected by TBI (Fig. 5m-r). The percentage of N-stage 2-3 cells in the SGZ was reduced (Fig. 5m, n), while the number in the GCL (GL1 and 2) was increased (Fig. 5m, o) in the TBI group. Similarly, the percentage of A-stage 1 cells in the SGZ was reduced (Fig. 5p, q), while their representation in Hilus was increased (Fig. 5p, r) in the TBI group. These results indicate a specific effect of TBI on the location of two NSC-derived cell populations in the DG, displacing them away from their native locations in the SGZ. Interestingly, Molecular Cartography showed an increase in GFAP+ cells in the hilus (Fig. 5s), in agreement with the general increase in GFAP levels observed in the hippocampus in the TBI group (Fig. 1a-d). Although all NSC-derived astrocytic cell populations express GFAP, A-stage 1 (and A-stage 4) cells are the two populations in which GFAP is expressed at high levels in the majority of cells (Fig. S3), suggesting that A-stage 1 cells contribute to this signal in the hilus.

**Figure 6.**
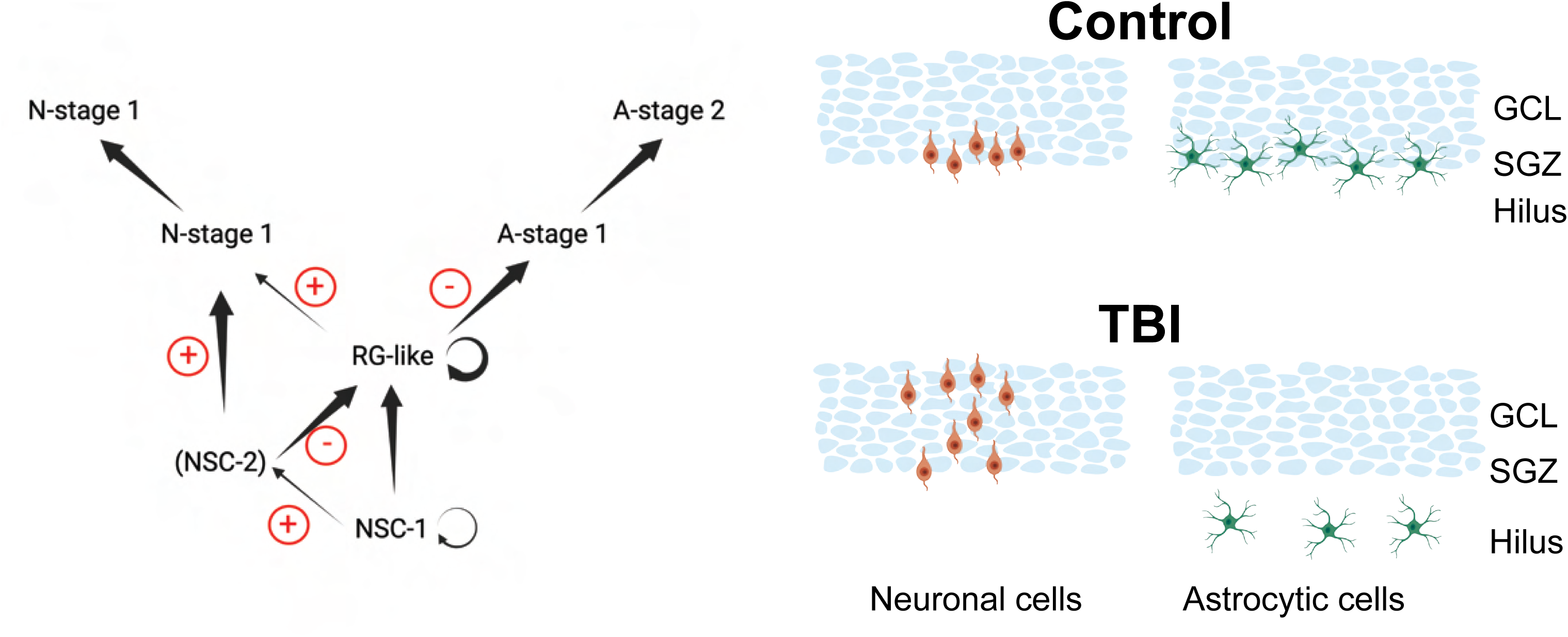
Schematic representation of the cell population-specific changes induced by TBI in NSC and NSC-derived cells in the DG. Schematic summary of cell transitions predicted by RNA velocity in the Control and TBI groups. Black arrows indicate the most probable transitions of NSCs and NSC-derived cells along neuronal and astrocytic lineages in uninjured AHN; arrow thickness indicates transition probability. Red +, transition promoted by TBI; Red-, transition inhibited by TBI; circular arrow indicates cell populations with predicted self-renewal potential (left). Schematic representations of GCL/SGZ/Hilus areas in the DG indicating the location of NSC-derived neuronal or astrocytic cells determined using Molecular Cartography spatial transcriptomics in both Control and TBI (right).

## Discussion

Under physiological conditions, native NSCs in the adult mouse DG generate new neurons and astrocytes, which integrate into the hippocampus’ complex cellular architecture and functional circuits, and contribute to hippocampal-dependent learning, memory and mood regulation (PMID: 31972145; PMID: 35451150, PMID: 33349709, PMID: 34103674). Here, we aimed to understand how TBI affects specific populations of NSC-derived cells in the DG. Using a combination of scRNA-seq, spatial transcriptomic and computational techniques, we: 1) identified distinct cellular populations in the AHN niche that belong to neuronal and astrocytic lineages; 2) observed that TBI induces changes in the relative proportion of cells in these lineages; 3) characterized changes in gene expression induced by TBI in specific NSC-derived cell populations; 4) confirmed the presence of NSC-derived neuronal and astrocytic cells in different subregions of the DG *in situ*; 5) identified a specific effect of TBI on the anatomical location of NSC-derived cell populations within the DG. Taken together, our results using a variety of methods, all support the conclusion that TBI modifies NSC fate to promote neurogenesis at the expense of astrogliogenesis, whilst also affecting the anatomical location of specific NSC-derived populations within the DG.

In agreement with observations made in other rodent TBI models, we observed an impairment in hippocampal dependent cognitive tasks 15 days after a controlled cortical impact, concomitant with hippocampal astrogliosis, indicated by increased GFAP expression and an increase in immature neurons (DCX+ cells) (PMID: 33622092, PMID: 10224295, PMID: 22900595, PMID: 33488351, PMID: 31670865). To perform the first unbiased assessment of the effects of mild-moderate TBI on the AHN, we performed single cell sequencing using cells isolated from Nestin-GFP mice, which express the fluorescent marker in NSCs, their progeny, and several other cell types within the niche (PMID: 14730584; PMID: 29241552). Using previously reported cell type markers, we were able to identify 10 cell clusters representing the major cell types expected in the DG: NSCs, neuronal and astrocytic cells, oligodendrocyte precursors and other less abundant cell populations, in agreement with previous studies (PMID: 29241552; PMID: 26299571). Following this first analysis, extraction of progenitors, astrocyte-like and neuron-like cells, with subsequent re-clustering and pseudotime-based lineage tracing, identified two specific cell lineages within the annotated cell populations (PMID: 32116127). Six neuronal cell populations (N-stage 1-6) fitted a lineage trajectory that initiated from three NSC populations (RG-like, NSC-stage 1 and 2), strongly suggesting they derive from them. An additional neuronal cell population (N-stage 7) did not fit within this lineage, suggesting a different origin, compatible with its lack of *Prox1* expression, a marker of granule cell identity in the DG (PMID: 22791897). Interestingly, four astrocytic cell populations (A-stage 1-4) fitted in a separate lineage trajectory, which also included RGL-like, NSC-1 and NSC-2 cells, indicating they also likely originate from these NSCs. Additionally, we found three astrocytic cell populations (A-stage 5-7) that did not fit within this trajectory, similarly suggesting a different origin. The three potential NSC populations we observed expressed *Ascl1, Vim, Hes5* and *Id4*, although the levels of these four markers were lower in NSC-stage 1 cells. Indeed, recent studies have identified populations of DG NSCs characterized by the differential expression of the pro-activation factor *Ascl1* and the inhibitor of DNA binding protein *Id4* (PMID: 31552825, PMID: 33349709, PMID: 33581058), indicating our NSC classification is consistent with earlier findings. This view is further reinforced both by Gene Ontology analysis, which confirmed that our NSC populations show high proliferative potential, and RNA velocity analysis which confirmed that the neuronal and astrocytic lineages described by our sequencing data are transcriptionally derived from these NSC populations. Independent confirmation of sequencing data was obtained using RNAscope *in situ* hybridization to detect NSC-derived cell populations. For our initial assessment, we focused on cells of the astrocytic lineage, as they represent a minority of NSC progeny, and are far less characterized in the literature than their neuronal counterparts (PMID: 31972145, PMID: 35451150). Specifically, we validated the presence of A-stage 1, 2 and 4 cells. We observed A-stage1 and 2 cells to be present mainly in the SGZ, an acknowledged anatomical location for NSCs and their immature progeny in physiological conditions (PMID:27814520, PMID: 35451150).

Crucially, all our data support the conclusion that TBI produces no change in the major cell types detectable in the AHN, or the developmental linages of NSC-derived neuron and astrocytes. This is in direct contrast to reports of unique cell populations associated with the development of Alzheimer’s (PMID: 32341542) and Huntington’s (PMID: 32070434) diseases, including the appearance of multiple populations of reactive astrocytes characterized by GFAP upregulation. Although these are chronic neurodegenerative diseases, in which changes in cell identity may well occur over years in a non-synchronous manner, large changes in cell identity were also reported on the shorter time-scale of EAE induction in mice (PMID: 29279367), suggesting that responses to injury and disease are a complex combination of initial cellular identity and insult (PMID: 32183137). Indeed, TBI-induced changes appear subtle, with the major effects appearing to be on the relative abundance of NSC-derived neuronal and astrocytic cells. We found that TBI resulted in an increase in the abundance of NSC-derived neuronal populations N-stage 1-5 cells. Concomitantly, TBI resulted in a decreased abundance of the NSC-derived astrocytic populations A-stage 1, 2 and 4, without seeming to have any effect on the abundance of A-stage 3 cells. These observations suggest a concerted effect of TBI on NSC-derived cell populations, possibly promoting neurogenesis at the cost of astrogliogenesis. Supporting this conclusion, RNA velocity analysis indicated that TBI altered the fate of RG-like cells, increasing their probability to transition to N-stage 1 cells and decreasing their probability to transition to A-stage 1 cells. Interestingly, NSC-stage 2 cells also showed an increased probability to transition to N-stage 1 cells, while NSC-stage 1 cells showed an increased probability to transition to NSC-stage 2 cells. These observations indicate an increase in neurogenesis in response to TBI. Further supporting this conclusion, N-stage 1 cells showed an increased probability of transitioning to more differentiated neuronal states – specifically N-stage 2 and 3. As N-stage 2 and 3 cells express neuroblast markers, this observation is compatible with an increase in immature neuronal cells induced by TBI in the DG, which we also observe using immunohistochemical detection of DCX. Overall, RNA velocity indicated an increased progression of progenitors along the neurogenic lineage at the apparent expense of astrogliogenesis following TBI. Interestingly, NSC proliferation and neurogenesis, induced by physiological stimuli such as running, are uncoupled from NSC-derived astrogliogenesis (PMID: 35451150). Therefore, the selective vulnerability of NSC-derived astrogliogenesis we report here may be related to the underlying pathology of TBI. Our analyses of DEGs in specific cell populations indicate that *Ppp1r14b* is upregulated following TBI in NSCs, astrocytic (A-stage 3), and neuronal (N-stage 1-4) cell populations. *Ppp1r14b* is a gene associated with proliferation and migration in other cell types (PMID: 36263632). *Ppp1r14b* is upregulated in hippocampal NSCs after kainic acid administration, an experimental condition that induces pathological NSC proliferation, and it is a validated target of microRNA-137, which prevents the kainic acid-induced loss of RG-like NSCs associated with hyperproliferation in the DG (PMID: 30837840, PMID: 25957904, PMID: 33642868). Taken together, these observations suggest that *Ppp1r14b*-mediated pathways may be involved in pathological activation and migration of NSCs and some of the populations that derive from them. However, the role of *Ppp1r14b* in hippocampal NSCs and their progeny remains unknown, and our observations warrant future studies to characterize its function(s) in detail. Changes in anatomical location of specific cell populations within the DG may be important to better understand the long-term consequences of TBI on hippocampal functions. Using Molecular Cartography and a multimodal reference mapping of our scRNA-seq dataset, we were able to localize 17 cell populations, that were initially annotated from the single cell RNA sequence dataset, to their location within the DG. Focusing on 13 populations, which represented NSCs and their progeny, we assigned cell populations to different anatomical locations in the AHN niche, and found that N-stage 3 and A-stage 1 cells, which represent two early cell populations from the NSC-derived neuronal and astrocytic lineages respectively, were misplaced in the DG of mice subjected to TBI. The alterations in the anatomical location of N-stage 3 is compatible with the increase in DCX+ cells observed with immunohistochemistry following TBI, and supported by our RNA velocity analysis. Changes in immature neuronal cell location impacts on their circuit integration, as they need to be physically adjacent to coordinate their lateral migration (PMID: 31346164). This suggests that N-stage 3 cells correspond to the ectopic neurons that have been observed after TBI in the DG (PMID: 26898165). Although previous studies have shown that astrocytic cells are generated in low numbers from NSCs in the AHN niche (PMID: 33349709, PMID: 35451150), molecularly distinct, spatially organized astrocyte populations appear to support local functions in the hippocampus ((PMID: 32139688, PMID: 36443610, PMID: 36378959, PMID: 36443610). Interestingly, recent observations have shown that physiological stimuli, such as running, stimulate adult NSC-derived neurogenesis without affecting astrogliogenesis (PMID: 35451150). In contrast, we show here that pathological insults, such as TBI, may stimulate adult NSC-derived neurogenesis at the expense of astrogliogenesis, highlighting the importance of assessing the balance of these processes in the AHN following pathological insults. In particular, our studies using RNA velocity and Molecular Cartography identified transition limited progression through A-stage 1 as a major effect of TBI on astrocyte maturation in the DG. Recent observations indicate that astrocytic cells are key regulators of cell-cell coordination in the hippocampus, supporting critical aspects of circuit function from synapse assembly and pruning, control of local homeostasis and modulation of synaptic transmission (PMID: 36748397). In particular, astrocyte dysfunction following TBI affects the metabolic support of neurons (PMID: 35951114) and hilar astrocytes contribute GABAergic inhibition of hippocampal dentate granule cells (PMID: 27161528). Moreover, neuronal rearrangements take place in the hilus after TBI, where mossy cells are sensitive and become hyperexcitable after (TBI PMID: 27466143; PMID: 10747187). Although it is tempting to speculate that the numerical and spatial changes in specific NSC-derived astrocytic populations may contribute to deficient metabolic support in the DG after TBI, future studies should clarify the functional relevance of these cell subpopulations. Overall, based on the results we describe, we propose a novel model for the effects of TBI on NSCs and the cell populations that derive from them in the AHN, which incorporates our key findings of changes in cell fate specification and differentiation and cell position (Figure 6).

Arguably, our work has some technical and conceptual limitations. Previous studies on the response of NSCs in the DG to TBI have delivered inconsistent observations regarding the degree to which neurogenesis is stimulated (PMID: 26414411). These discrepancies may be explained by variations in the use of different TBI models and experimental design. A previous study using the CCI model to investigate the effect of injury severity, concluded that moderate TBI promoted NSC proliferation without increasing neurogenesis, as measured by the number of DCX+ cells in the DG two weeks after injury (PMID: 26414411). In contrast, we observed a significant increase in DCX+ cells, although we used different injury induction parameters. According to a recent effort to standardise CCI parameters across different laboratories, the TBI that we induced can be defined as mild to moderate (PMID: 30017882). This definition is consistent with our observation of an impairment in hippocampal-dependent learning in the Morris Water Maze (a cognitive test frequently used to evaluate mild to moderate TBI), concomitant with an increase in the number of DCX+ cells (PMID: 30017882). Regarding possible implications for TBI in humans, CCI only mimics certain aspects of human TBI (PMID: 30017882). Specifically, mild, moderate, or severe TBI, are clinically defined in humans based on loss of consciousness, alterations in mental states, post-traumatic amnesia or coma at different times after TBI, some of which are not considered in rodent models (PMID: 22834016; PMID: 30017882).

While the presence of adult hippocampal neurogenesis in humans has been recently debated (PMID: 29681514; PMID: 31899214), particularly in respect of how its impairment in older adults might play a role in neurodegenerative disease (PMID: 34672693; PMID: 35420933; PMID: 35420939; PMID: 35420954), most studies agree on its occurrence in younger individuals (PMID: 29513649) and a possible a roadmap toward a better understanding of the role of AHN in AD, for which scRNA-seq studies may be crucial, has been recently proposed (PMID: 36736288). Importantly, TBI induces the expression of NSC markers in individual cells of the perilesional human cortex, suggesting that TBI induces neurogenesis in the human brain (PMID: 21275797).

In conclusion, therefore, the molecular and cellular changes we describe here in mouse may well help us to better understand the changes induced by TBI in the hippocampus of young TBI patients, possibly opening up new targets for therapeutics.

## Methods

### Animals

Eight-week-old male Nestin-GFP^+/-^ or C57/Bl6J (wild type) male mice were used in our experiments. All mice were bred in house and housed in groups of 3-4 animals per cage throughout the experiment under a 12-h light/dark cycle (lights on at 08.00AM) in a temperature and humidity-controlled room (21°C, 50%) with *ad libitum* access to food and water. All animal procedures were approved by the commission for Animal Welfare at the University of Amsterdam (CCD 4925) and/or VIB-KU Leuven (082/2018) and were performed according to the guidelines and regulations of the European Union for the use of animals for scientific purposes and the ARRIVE guidelines for reporting animal research (PMID: 32663219). All mice were randomly assigned to experimental groups.

### Controlled cortical impact

A controlled cortical impact model was used to induce a traumatic brain injury of mild to moderate severity, as previously described (PMID: 35618831). In brief, mice were placed in a stereotaxic frame and anesthetized using 2% isoflurane throughout the surgery. A craniotomy was performed, creating a window along the skull sutures from bregma to lambda over the left hemisphere. The impact piston was placed at an approximately 20-degree angle directly on the brain surface and an impact was performed using the following settings: 1 mm impact depth, 5.50 m/s velocity, dwell time 300 ms. After the impact, the skull bone was placed back and glued in place using Superglue. The skin was stitched to close the wound and animals were allowed to recover on a 37°C heat pad until fully awake.

### Tissue extraction

Mice were sacrificed 15 days post-TBI. Mice destined for immunohistochemistry were sacrificed by an overdose of Euthasol, followed by intracardial perfusion-fixation with ice-cold PBS, followed by 4% PFA. The brains were isolated and stored in PBS until further use. Mice destined for single cell experiments were sacrificed by rapid decapitation, after which brains were removed and the dentate gyri microdissected ready for dissociation. Mice destined for experiments using either RNAscope or Molecular Cartography were sacrificed by rapid decapitation, after which the brains were removed and directly processed, according to the manufacturer’s protocols.

### Immunohistochemistry

PFA-fixed brains were cryoprotected using 30% sucrose and then sliced into 40 μm-thick slices: every 8^th^ section was taken for immunostaining, ensuring a 280 nm separation between slices, as previously described (PMID: 31222184). Fluorescence immunohistochemistry was performed following a standard procedure. Sections were first incubated with blocking and permeabilization buffer (1X PBS / 5% normal goat serum (Cell Signaling, cat #5425) / 0.3% Triton X-100) for 30 minutes, followed by incubation with the primary antibody overnight at 4°C. Sections were thoroughly washed with PBS and subsequently incubated with fluorescent secondary antibodies for 2 hours at room temperature. After thorough washing with PBS, tissue slices were mounted on glass slides and counterstained with Vectashield antifade mounting medium containing DAPI (Vector Laboratories, cat# H-1200-10). The following antibodies were used: rabbit anti-GFAP (1:10,000 dilution; DAKO, cat# Z0334) in combination with goat anti-rabbit Alexa 568 (1:500 dilution; Thermo Fisher Scientific/Invitrogen, cat # A-11011), goat anti-DCX (1:500 dilution; Santa Cruz, cat# sc-8066) in combination with donkey anti-goat Alexa 488 (1:500 dilution; Thermo Fisher/Invitrogen, cat # A-11055). DAB-based immunohistochemistry for DCX was performed according to a standard protocol, as previously described (PMID: 22925833), using goat-anti DCX (1:800 dilution; Santa Cruz, cat# sc-8066) in combination with donkey anti-goat-biotin (1:500 dilution; Jackson Immuno Research, cat# 705-065-147). Nissl staining was performed to counterstain nuclei using Cresyl Violet, as previously described (PMID: 31222184). Total DCX+ cells were assessed using a stereological approach, as previously described (Schouten et al., 2015). Gliosis was analyzed by measuring GFAP surface area coverage in the whole hippocampus using ImageJ as described before (PMID: 30837840).

### RNAscope® fluorescent multiplex *in situ* hybridization

Freshly removed brains from C57/Bl6J mice were directly snap-frozen using liquid nitrogen. 16 µm-thick sections containing the hippocampus were cut using a cryostat and collected directly on Superfrost glass slides. RNA transcripts of interest (*Slc1a3, Hapln1, Neat1, Sned1, Sparc, Ascl1, Frzb*) were targeted using RNAscope^®^ probes. The RNAscope^®^ HiPlex v2 Assay was performed following the manufacturer’s instructions as described in (PMID: 22166544), and images were collected on a Zeiss LSM 510 confocal laser-scanning microscope (10x air, 40x water objectives). A 10x overview of the dorsal DG was taken first and 40x Z-stack images with 1 µm intervals were then produced and analyzed using ImageJ. The SGZ was defined as two cell bodies distance from the GCL (PMID: 16267214).

### Molecular Cartography

C57/Bl6J mouse brains were processed according to the manufacturer’s protocol. Briefly, the ipsilateral side of the brain was trimmed to a dimension of maximum 1 cm thickness and immersed in a proprietary fixative (Resolve BioSciences GmbH, Monheim am Rhein, Germany), after which brains were placed in a proprietary stabilization buffer. Brains blocks were then sectioned into 2 mm slices and immersed in cryo-embedding medium (Resolve BioSciences GmbH, Monheim am Rhein, Germany), followed by snap-freezing in liquid N_2_. 10 µm sections were generated using a cryostat. Samples processed in this way were used for highly multiplexed single molecule *in situ* hybridization (Molecular Cartography platform) as described in (PMID: 35021063). Essentially, five tissue sections from three different animals (TBI) and four tissue sections from two different animals (control) were stained with probes targeting 93 genes defining specific stages along both the neuronal and astrocyte lineages (Table S9). Tissue sections were counter-stained with DAPI to allow cell identification through nuclear segmentation. 9 tissue sections were imaged and data files generated containing the x-y co-ordinates of each transcript detected. These images were processed by QuPath 0.2.3 software to segment single cells from the granule layer of the dentate gyrus (DG) (sub-divided in GL1 and GL2), subgranular zone (SGZ) and Hilus (PMID: 30837840), based on the DAPI signal (using QuPath’s cell detection algorithm with the parameters indicated in Table S10). Co-ordinates for segmented nuclei were transferred to ImageJ 2.0.0-rc-43/1.52n and the transcript count per nucleus was extracted using the Polylux_V1.6.1 ImageJ plugin, developed by Resolve Biosciences.

### Single Cell Suspension

A single cell suspension was made from the micro-dissected dentate gyri using a neural tissue dissociation kit (Miltenyi Biotech), according to the manufacturer’s protocol. In brief, 7 ipsilateral dentate gyri were collected per experimental condition. Dissociation enzymes were added followed by manual dissociation using first a P1000 pipet tip, followed by further dissociation using a P200 pipet tip. After full dissociation, the suspension was filtered using a 40 µm cell strainer and collected in HBSS containing RNase inhibitor. The suspension was then centrifuged at 300xg_Av_ for 12 minutes, after which the cell pellet was resuspended in PBS containing 0.5% FBS.

### Fluorescence activated cell sorting

GFP positive cells were isolated using fluorescence activated cell sorting (FACS) (BD FACSAria™ III Cell Sorter), as recently described (PMID: 31222184). Propidium iodide (PI) was added to the single cell suspension to discriminate between live and dead cells. In a first step, doublets and cell debris were removed based on forward and side scatter. Dead cells were then removed based on PI staining, after which the single cell suspension was sorted into GFP positive and negative populations based on the intrinsic GFP signal recorded in the FITC channel. GFP positive populations were sorted and collected in PBS containing FBS. Cell concentration was adjusted to 1000 cells/µL.

### Library preparations

Single cell suspensions were prepared from microdissected DGs obtained from 8 weeks old Nestin-GFP mice, using 5-6 animals per condition (Control and TBI). Library preparations for scRNA-seq were performed using a 10X Genomics Chromium Single Cell 3’ Kit, v3 (10X Genomics, Pleasanton, CA, USA). The cell count and the viability of the samples were assessed using a LUNA dual florescence cell counter (Logos Biosystems) and a targeted cell recovery of 6,000 cells per sample was aimed for. Post-cell counting and QC, the samples were immediately loaded onto a Chromium Controller. Single cell RNAseq libraries were prepared according to the manufacturer’s recommendd protocol (Single cell 3’ reagent kits v3 user guide; CG00052 Rev B), with library quality checked at all indicated protocol points using a Qubit to measure cDNA concentration (ThermoFisher) and Bioanalyzer (Agilent) to measure cDNA quality. Single cell libraries were sequenced on either an Illumina NovaSeq 6000 or HiSeq4000 platform using a paired-end sequencing workflow with the recommended 10X v3 read parameters (28-8-0-91 cycles). We aimed for a sequencing coverage of 50,000 reads per cell.

### SC data preprocessing and clustering analysis

Data were demultiplexed and mapped using a standard CellRanger 3.0.2 workflow making use of the UCSC mouse genome GRCm38/mm10 assembly and refdata-cellranger-mm10-3.0.0 reference dataset. Libraries having low or high RNA content were removed to exclude cells with degraded RNA or potential cell doublets, respectively. Clustering analysis was done using the R Seurat_3.2.0 package (PMID: 31178118) using the standard workflow. *Malat1* was excluded from the analysis, since many reads mapped to it, creating an artificial peak. Datasets were normalized by the global-scaling normalization function *LogNormalize*. Variable features were found using the *FindVariableFeatures* default variance stabilizing transformation (vst) method, by fitting log(variance) and log(mean) using local polynomial regression (loess). Canonical Correlation Analysis (CCA) was performed for the integration of anchors by *FindIntegrationAnchors* on 20 CCA dimensions and *IntegrateData* on the top 20 CCAs. The integrated data was scaled and centered using the *ScaleData* function on all transcripts. Principal Component Analysis (PCA) on the resulting data was performed using 30 dimensions. A resampling *Jackstrow* test was performed to assess the significance of PCA components. The percentage of variance explained by each PCA was saturating at number 20. Therefore, the 20 most significant PCs were selected for UMAP reduction using *RunUMAP*, *FindNeighbors* and *FindClusters* functions. Except for the number of PCA components, no other default parameters were changed in these functions. Sub-clustering was done using the same procedure, except that only cells belonging to the astrocytic and neuronal lineages (including NSC and progenitor cells) were used (taking into account the top 30 CCA and top 16 PCA dimensions). The resolution of the initial high-level cell type clustering was set to 3.0, and when reclustering data it was set to 0.8.

### Population enrichment analysis

The significance of population enrichment by TBI or Control samples was tested with a 2-sided binomial test, using the R base *binom.test* function. The probability of success was set to represent the proportion of sequenced cells originating from the TBI samples against the whole database (62.96% for the higher order clustering (Fig. 2c) and 57.39% in case of the pseudotime analysis (Fig. 2f).

### Differential expression analysis

After re-clustering, astrocyte and neuron data went through an additional round of SCT normalization, performed using the Seurat *SCTransform* function with default parameters, to regress out noise arising from mitochondrial genes. Differential gene expression analysis to detect cell-type specific markers (see Table S3) was performed on the SCT normalized data using the Seurat *FindMarkers* function, using a 0.25 threshold on the ln-fold expression difference and requiring a minimum of 25% of the cells in a given population to express the marker gene. The significance of each marker was calculated using a Wilcoxon Rank Sum test and corrected using the Bonferroni method. To detect differentially expressed genes between equivalent cell populations identified in control and TBI conditions, MAST tests were performed (using an extension in the Seurat *FindMarkers* function) with subsequent Bonferroni correction (Table 6). Genes were taken to be differentially expressed (DEGs) if they showed at least a 0.2 ln-fold change in a minimum of 25% of cells for a given population

### Integration of the scRNA-seq and spatial data

Spatial and single cell data were integrated in order to find matching populations. For this purpose, all the extracted spatial data was merged with the single cell data, using the R Seurat_3.2.0 package anchor-integration method. Prior to integration, spatial and single cell datasets were separately log-normalized and variable features for each were identified, using *LogNormalize* and *FindVariableFeatures* with vst methods, respectively. Next, integration anchors were found using the *FindIntegrationAnchors* function, using 8 dimensions of CCA reduction. Next, the two databases were merged using the *IntegrateData* function, using 8 CCA dimensions. Finally, the integrated data was scaled; PCA and PCA-based UMAP analysis were performed on 8 PCA dimensions using *RunPCA*, *RunUMAP*, *FindNeighbors* and *FindClusters* functions with resolution set to 1 and all other parameters set to default. Clouds representing only the spatial data were identified. These clusters are either driven by technical factors or represent mature cells (not present in the single cell database) and were, therefore, removed before the integration method described above was repeated using the following parameters: 10 CCA, 10 PCA and 0.8 clustering resolution.

### Pseudotime analysis

Lineages were constructed using the Monocle v3_1.0.0 R package (PMID: 30787437), using the *cluster_cells, learn_graph* and *order_cells* functions with default parameters, based on clustering obtained using the UMAP dimensionality reduction method and the integrated normalized expression matrix in Seurat. NSC-like populations (NSC-stage1, NSC-stage2, RG-like for the 10x scRNA dataset and clusters 7-8 (Fig. 5K) for the integrated spatial and scRNA datasets) were used as the root from which developmental pathways were developed. N-stage 7 cells and A-stage 5-7 cells clustered separately in UMAP space, indicating that they likely represent non-NSC-derived lineages. Therefore, they were excluded from pseudotime analysis.

### Pathway analysis

Gene-enrichment and functional annotation analyses of cell population-specific differentially expressed (up/down regulated) genes were performed using GO and KEGG databases accessed through clusterProfiler version 3.18.1 (pmid 34557778). First, an unbiased analysis was performed to report all the KEGG pathways/GO terms related to the gene list of interest. This was followed by sorting based on the following keywords: *’Synapse’, ‘Axon’, ‘Neuron’, ‘Nervous’, ‘Glia’, ‘Astrocyte’, ‘Microglia’, ‘Injury’, ‘Growth’, ‘Tnf’, ‘Neuro’, ‘Age’, ‘Myelin’, ‘Sheath’, ‘Reactive’, ‘Ion’, ‘Proliferation’, ‘Genesis’, ‘Development’, ‘Morphology’, ‘Formation’, ‘Circuit’, ‘Axonogenesis’*. The corresponding outputs for the biased and unbiased analysis are reported in Tables S4, S5 (for marker genes) and Tables S7, S8 (for up/down regulated genes in TBI).

### RNA Velocity Analysis

Analysis was performed on the R velocyto.R_0.6 package, using spliced and unspliced RNA counts obtained through the standard run10x workflow using the gRCm38/mm10 genome. UMAP embedding space was imported from the respective Seurat clustering analysis. RNA velocity was calculated for control and TBI separately, after removing the N-stage 7 and A-stage 5,6 and 7 populations. The following workflow was implemented:

#### Step1

*RunVelocity* function was used with default parameters, except *spliced.average = 0.5, fit.quantile = 0.02, kCells = 20* to obtain “current” and “deltaE” matrices.

#### Step2

The “current” and “deltaE” matrices from step1, and the UMAP embeddings from clustering analysis, were used in the *show.velocity.on.embedding.cor* function to obtain “tp” (transition probability matrix), “cc” (velocity-correlation matrix) and RNA Velocity on embeddings. All parameters were used with default settings, except the neighborhood size which was set to *n=300*.

#### Step3

For every population, the mean transition probability to each of its neighbors was calculated, using the *show.velocity.on.embedding.cor* function, where *emb* represented the UMAP embeddings of the clustering analysis (Fig. 2d) and *vel* represented the list of “current” and “deltaE” matrices (step1) for the subpopulation under consideration. The neighborhood size, *n,* was set according to the number of cells in the populations of interest, using a minimum of 300 cells. The velocity-correlation matrix, *cc*, was obtained from step2 for the population of interest. Parameter *scale=’sqrt’* was used. The remaining parameters were used with default settings. The “tp” matrices obtained after step3 were used in step 4.

#### Step4

Using “tp” matrices obtained from step3, barplots were constructed. Error bars show the average ± SEM. * marks a significant difference between control and TBI, based on independent non-parametric Wilcoxon tests. (ns: p > 0.05, *: p <= 0.05, **: p <= 0.01, ***: p <= 0.001, ****: p <= 0.0001). Codes are available at https://github.com/araboapresyan/rna_velocity_analysis.

### Figure preparation

Figures were prepared using R v3.6.0/v4.1.0, RStudio 1.0.136/1.4.1106/v4.1.2, Adobe Photoshop CC 2022, Adobe Illustrator 2022 and Graphpad Prism 9.

## Acknowledgements

PB, GM, NR, WK, RAC, IvdV, ET, ID, JME and CPF were supported by the European Union’s Horizon 2020 research and innovation program ERA-NET-NEURON (grant EJTC 2016) to CPF and JME, the Netherlands Organization for Scientific research (NWO), and Alzheimer Nederland. PJL is supported by the Center for Urban Mental Health and by Alzheimer Nederland. MGH’s work in Leuven was supported by the Belgian Scientific Research Fund (Fonds Wetenschappelijk Onderzoek – FWO - Grants G066715N, 1523014N and I001818N) and the Belgian Alzheimer’s Society (SAO) (Grant S#16025). MGH is currently the ERA Chair (NCBio) at i3S Porto, funded by the European Commission (H2020-WIDESPREAD-2018-2020-6; NCBio; 951923). AM acknowledges funding from the Stichting Alzheimer Onderzoek (SAO #2020034) and VIB Tec Watch funding. FP was supported by a Fundação para a Ciência e a Tecnologia (FCT) Ph.D. fellowship (2020.08750.BD).

## Author contributions

PB designed and performed all the in vivo animal experiments, including controlled cortical impact, behavior, tissue extraction, sample preparation for transcriptomics, immunohistochemistry and RNAscope, assisted by GM, NR, WK, RAC, IvdV, ET and ID, analyzed and interpreted the data derived from these experiments, composed the first draft of the manuscript and figures and supported AM in computational analysis. PJL contributed to experimental design, provided funding and corrected the manuscript, JME supervised optimization of the controlled cortical impact technique, contributed to experimental design conception of the experiments, data interpretation and corrected the manuscript. AM performed all computational analysis, with the assistance of SH, except for the RNA velocity analysis, which was performed by AA. AM prepared relevant figures and provided input on the manuscript. FP processed samples for Molecular Cartography experiments. SP supervised tissue dissociation, sequencing library preparation and sequencing. MGH provided funding, access to the single cell sequencing facility at VIB-KU Leuven and arranged early access to the Molecular Cartography platform. MGH supervised data acquisition, computational analysis and interpretation (with input from TGB) and redrafted the manuscript. BN, AB and NK performed Molecular Cartography experiments and provided input on data analysis. CPF conceived the experiments, provided funding participated in experimental design, supervised experiments and data interpretation, composed the final version of the figures and wrote the final version of the manuscript in consultation with MGH.

## Conflict of interest

MGH has acted as a paid consultant to Resolve Bioscience during development of their Molecular Cartography platform.

## Figure legends

**Figure S1.**
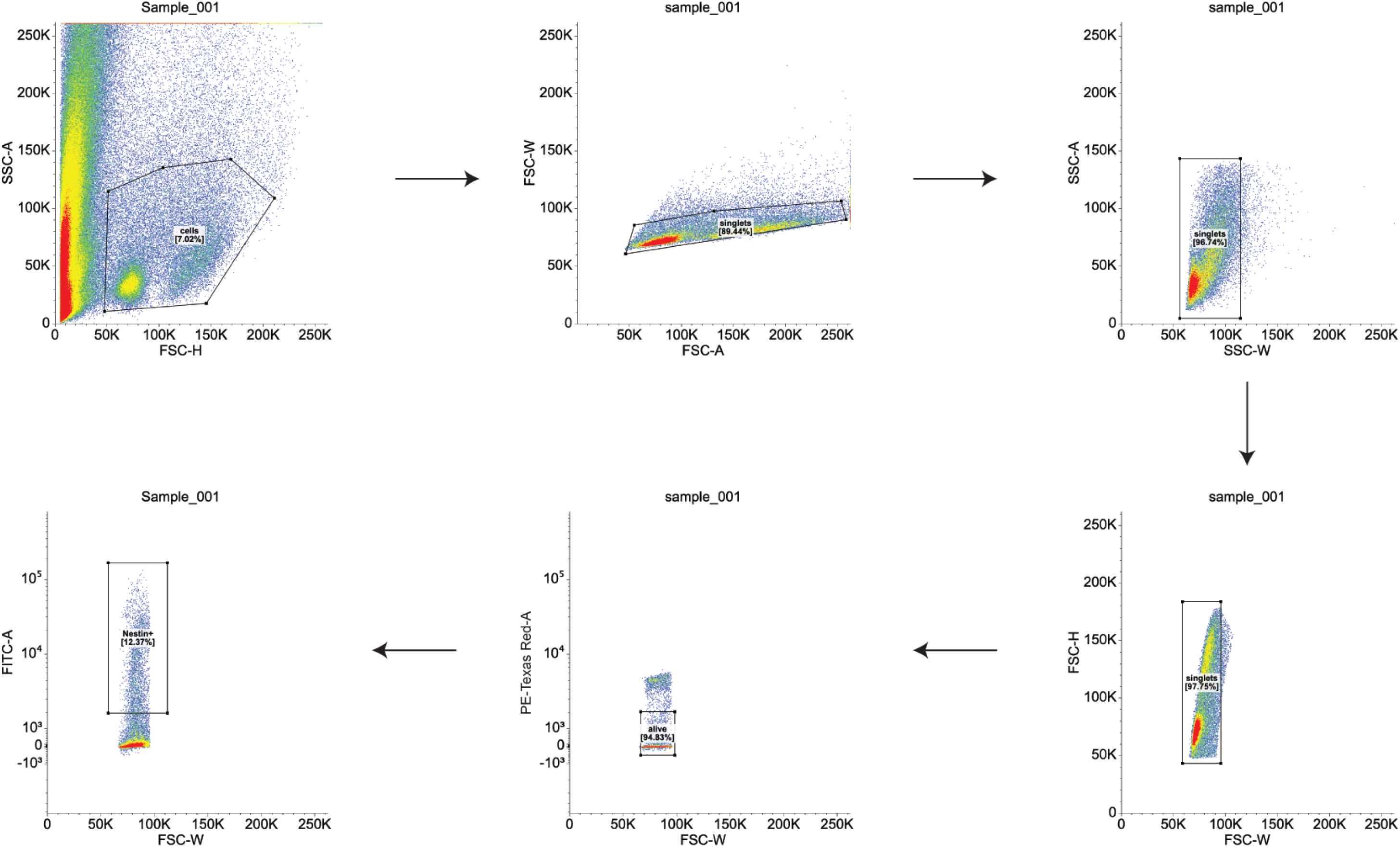
Gating strategy used for FACS-based purification of GFP+ cells. Cells were initially selected using Forward and Side Scatter area plots (FSC-A and SSC-A) (indicated gate). To minimize the amount of small debris collected, care was taken to adjust the lower limit on the forward scatter (measure of size) axis, although the gate was left wide enough that smaller cells were still captured. Cell doublets were excluded using forward/side scatter width vs height plots (FSC-W/FSC-H and SSC-W/SSC-H). Live-dead staining using Propidium Iodide was used to identify viable cells, which were subsequently separated into GFP+ and GFP- populations based on fluorescence. Nestin+ cells were retained for further use.

**Figure S2.**
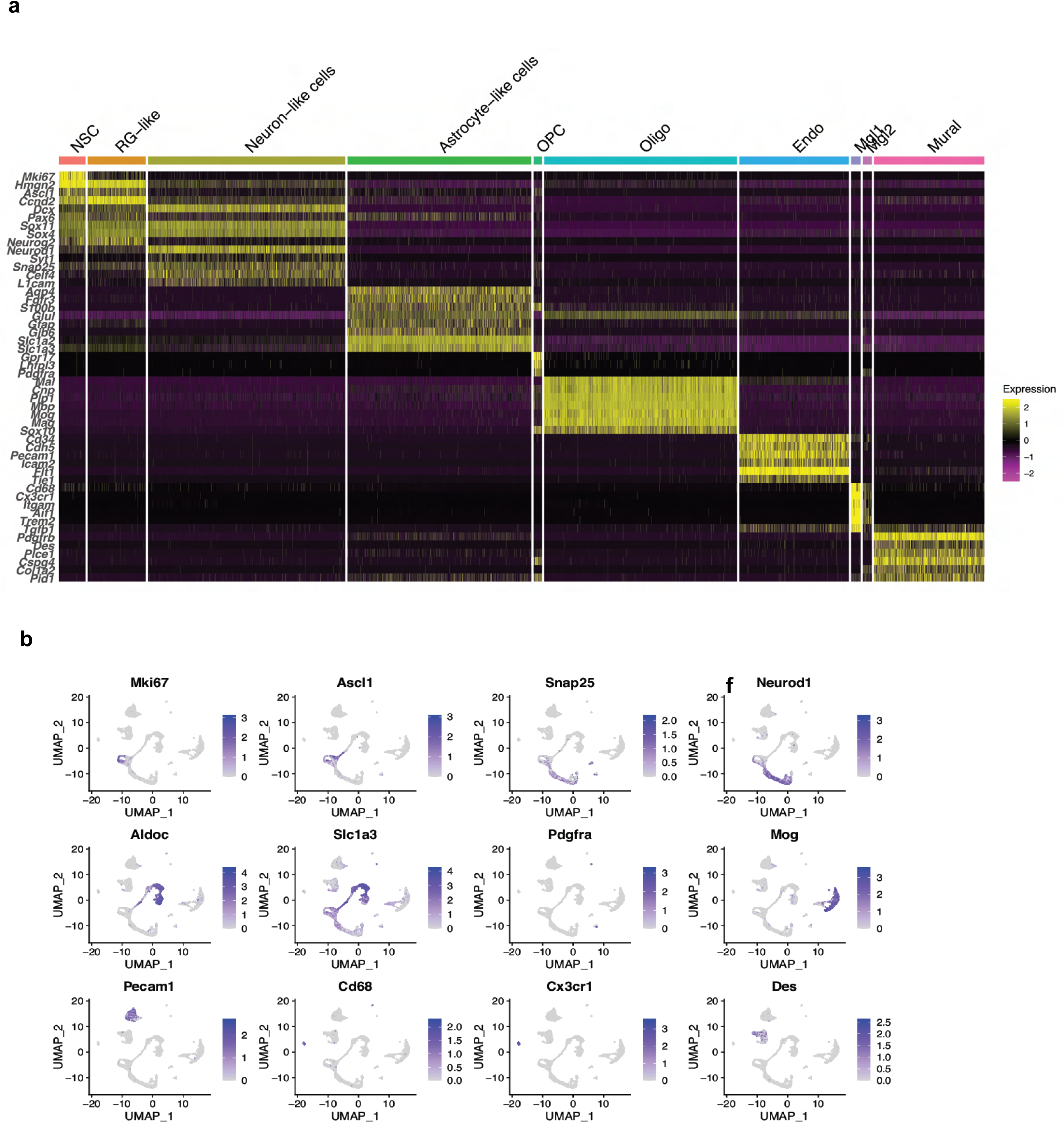
Cell type assignment based on specific marker gene expression. Gene expression heatmap for higher-order cell types (columns) grouped according to the Seurat classification shown in Fig. 2a. Color-coding from Fig. 2a is retained. Magenta, low expression; yellow, high expression, ln-normalized gene expression data is shown. UMAP representations showing expression patterns of indicated marker genes across cell clusters (b), color bars indicate relative intensity of expression, SCT normalized values are shown.

**Figure S3.**
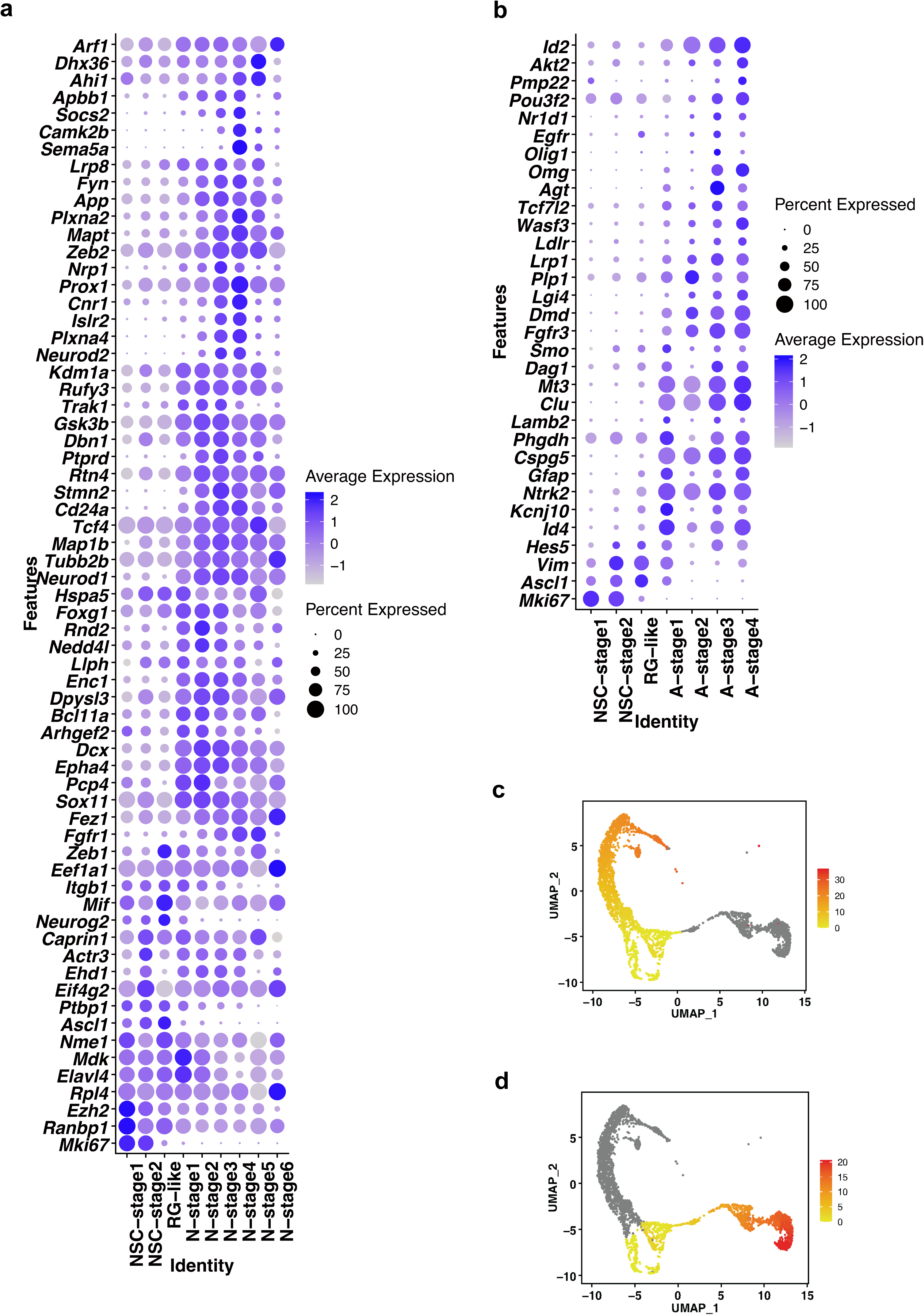
Pseudotime assignment of NSC-derived cell lineages defined by expression of specific marker genes. Dot-plot graphs indicating the relative expression of marker genes in NSCs and NSC-derived neuronal (a) and astrocytic (b) populations; scaled SCT normalized data are shown. Color bars represent relative intensity of gene expression, dot size represents percentage of cells within an individual cell cluster (identity) expressing the indicated marker gene (feature). UMAP representations showing NSC-derived neuronal (c) and astrocytic lineages (d); color bars indicate calculated pseudotime distances from NSCs (yellow: minimum, red: maximum).

**Figure S4.**
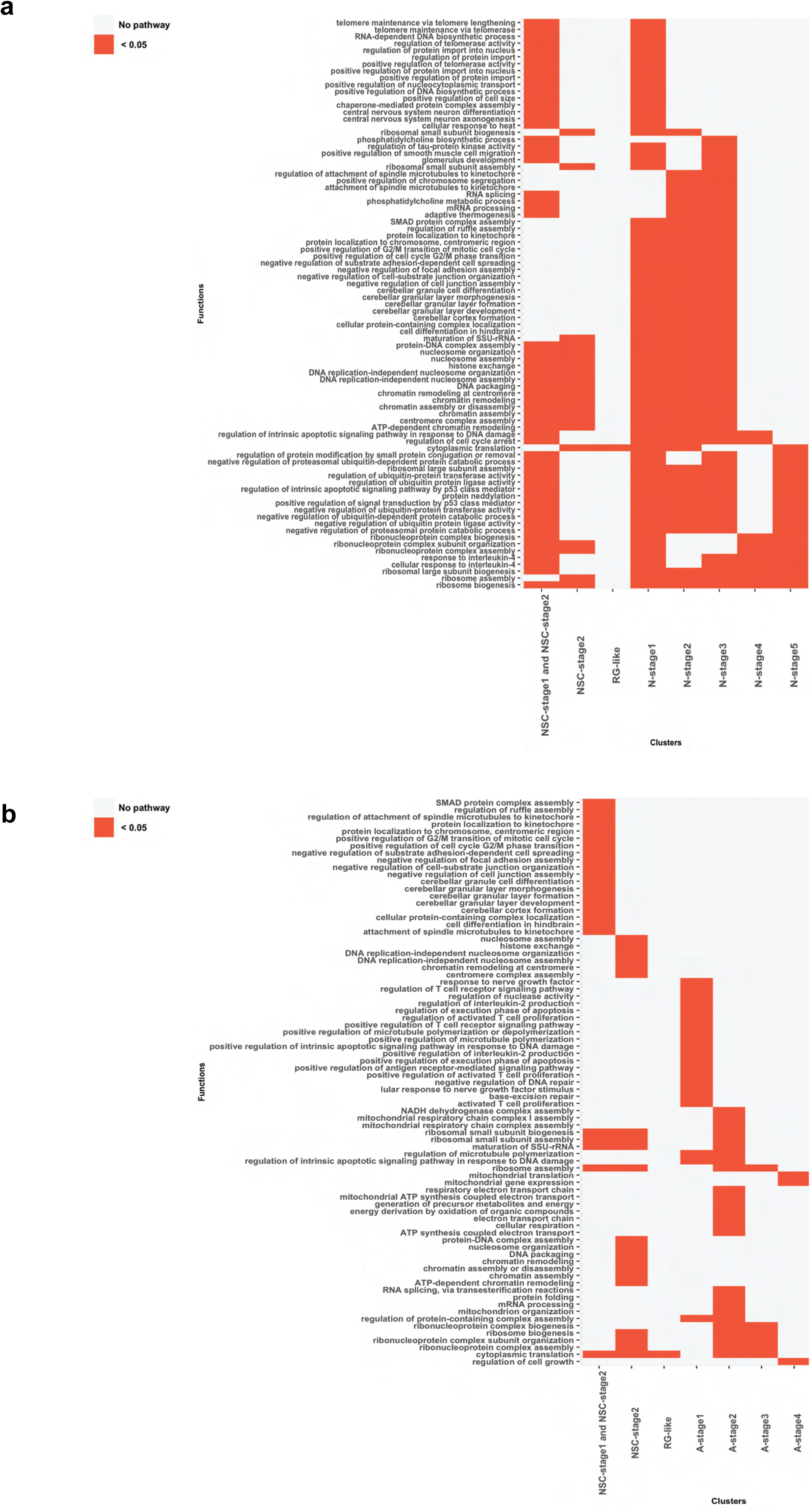
GO biological pathway matrix for the differentially expressed genes in NSC-derived cell populations. Differentially over-represented (red, p<0,05) biological pathways found in the neuronal (a) and astrocytic (b) lineages.

**Figure S5.**
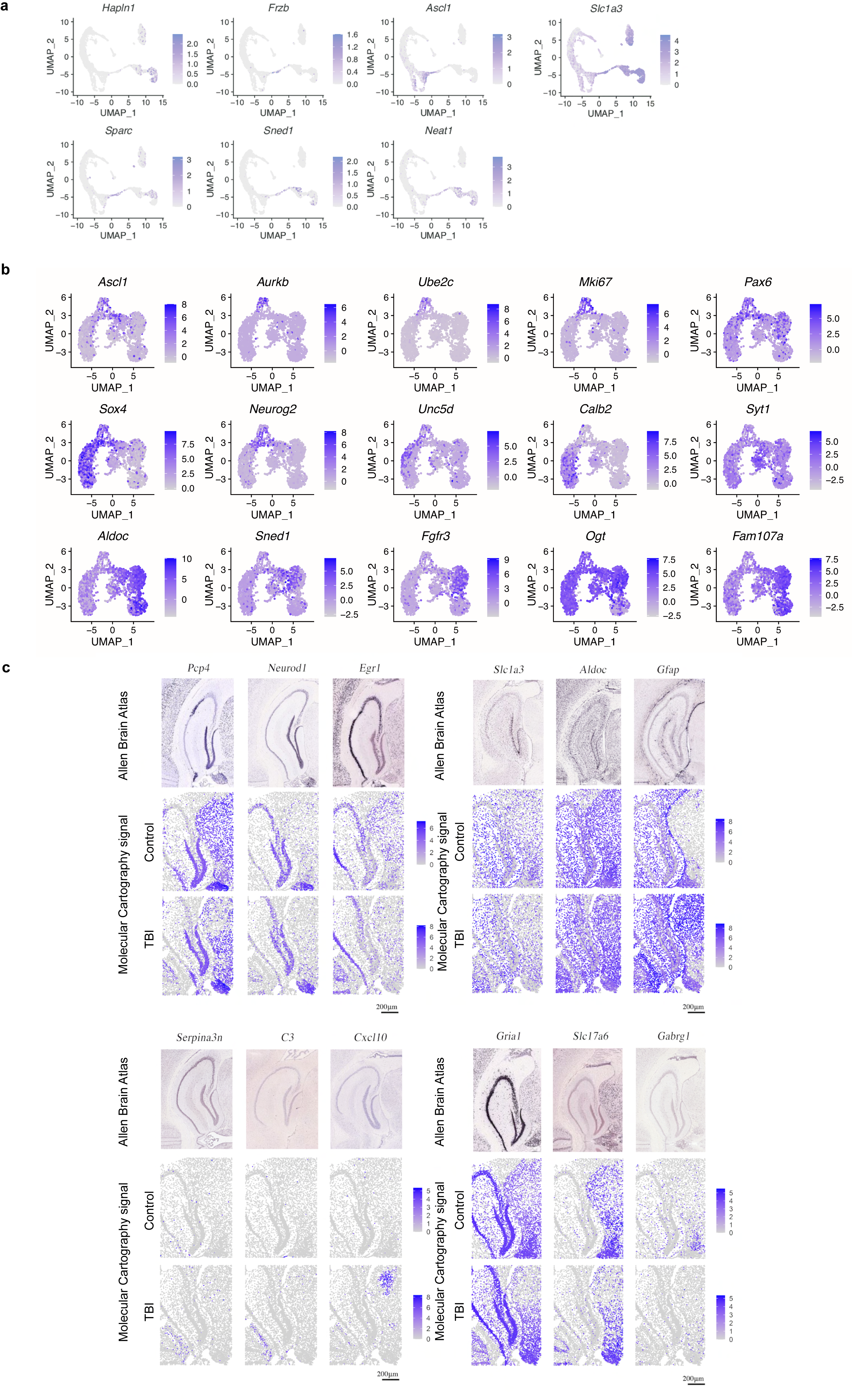
Design and validation of probes for spatial transcriptomics. UMAP plots showing the expression of individual gene markers used for RNAscope validation, across astrocytic cell clusters, SCT normalized values are shown (a). UMAP representations of the combined 10X and Molecular Cartography dataset, indicating expression of exemplar marker genes for the various identified cell populations (b). Example images showing how mRNA expression detected by the 12 Molecular Cartography probes indicated at the top of each panel compares to that reported in the Allen Brain Atlas (c), colored bars indicate relative intensity of gene expression, raw counts are shown. Colored dots indicate individual cells in the dentate gyrus.

